# Interaction between habits as action sequences and goal-directed behavior under time pressure

**DOI:** 10.1101/2022.11.15.516603

**Authors:** Sascha Frölich, Marlon Esmeyer, Tanja Endrass, Michael N. Smolka, Stefan J. Kiebel

## Abstract

Human behaviour consists in large parts of action sequences that are often repeated in mostly the same way. Through extensive repetition, sequential responses become automatic or habitual, but our environment often confronts us with events to which we have to react flexibly and in a goal-directed manner. To assess how implicitly learned action sequences interfere with goal-directed control, we developed a novel behavioural paradigm in which we combined action sequence learning through repetition with a goal-directed task component. So-called dual-target trials require the goal-directed selection of the response with the highest reward-probability in a fast succession of trials with short response deadlines. Importantly, the response primed by the learned action sequence is sometimes different from that required by the goal-directed task. As expected, we found that participants learned the action sequence through repetition, as evidenced by reduced reaction times and error rates, while still acting in a goal-directed manner in dual target trials. Specifically, we found that the learned action sequence biased choices in the goal-directed task towards the sequential response, and this effect was more pronounced the better individuals had learned the sequence. Our novel task may help shed light on the acquisition of automatic behavioural patterns and habits through extensive repetition, allows to assess positive features of habitual behaviour (e.g. increased response speed and reduced error rates), and importantly also the interaction of habitual and goal-directed behaviours under time pressure.

## 1 INTRODUCTION

Our daily lives are governed by habitual behaviour patterns (Wood et al., 2002; James et al., 1890). For example, most people will use the same road to get to work every day without giving it much thought. At the same time, in our dynamic environment we cannot solely rely on habitual behaviour because we have to be able to flexibly respond in a goal-directed manner to events that require reevaluation of the situation and the comparison of different response options. To stay with the above example, on the way to work we would usually take the same route but we must be able to quickly switch away from our habitual behaviour to a goal-directed reappraisal of the situation, as in the case of an accident that blocks our way to work.

The terms habit and automaticity are sometimes used as synonyms, although these two concepts are clearly not identical, albeit closely related (Garr and Delamater, 2019; Wood et al., 2014; Hardwick et al., 2019). Without going into too much detail here, habits can be considered as a kind of automatic, implicit process, but different from other automatic processes like for instance priming, Pavlovian conditioning, or automatic goal pursuit (Mazar and Wood, 2018; Moors and De Houwer, 2006). Habitual and automatic behaviour have been explained with dual-process theories, suggesting that the human brain comprises two different processing modes, an automatic, unconscious and fast processing system, sometimes called *system 1*, and a slow, conscious, and deliberative system, called *system 2* (Wood et al., 2014; St Evans et al., 2008), along with an arbitrator system which determines the influence of those two systems. In this framework, habitual and automatic behaviours belong to system 1, as opposed to goal-directed behaviour, which is attributed to system 2. The application of the dual-process theory to human behaviour aligns with a broader field in behavioural and cognitive sciences, where not only behaviour but also higher human cognition and reasoning is described in terms of the dual-process theory (Evans and Stanovich, 2013; St Evans et al., 2008; De Neys and Pennycook, 2019; Bellini-Leite, 2022; Milli et al., 2021; Grayot, 2020). While the validity of the dual-process theory has repeatedly been put into question (Kruglanski and Gigerenzer, 2018; Osman, 2004), a wealth of evidence indicates that automatic/habitual behaviour and goal-directed behaviour involve two different neural pathways (Yin and Knowlton, 2006; Graybiel et al., 2008; Dolan and Dayan, 2013). More specifically, studies in humans and animals have shown that goal-directed behaviour is executed under the involvement of cortico-striatal loops between the basal ganglia and parts of the prefrontal cortex, while automatic and habitual behaviours involve a distinct loop between the basal ganglia and the sensorimotor cortex (Yin and Knowlton, 2006; Graybiel et al., 2008; Dolan and Dayan, 2013).

An influential approach for testing the relative influences of the goal-directed and the habitual/automatic systems in human decision making is the two-stage task. Behavioural results have been modelled as the balance between a model-based (MB) reinforcement learning controller and a model-free (MF) controller (Daw et al., 2005, 2011). However, action-outcome associations in the original two-stage task change over time, which puts into question its capacity to induce habits. Furthermore, while the MB controller seems to reliably model goal-directed behaviour, the MF controller’s influence on behaviour has been reported not to correlate with measures of habitual behaviour (Gillan et al., 2015; Sjoerds et al., 2016). Furthermore, model-free/model-based (MF/MB) reinforcement learning models as used for the analysis of the probabilistic two-stage task with dynamic reward probability (Daw et al., 2011) typically infer the interplay between the MF and MB controllers as a smoothed estimator based on the recent trial history. This smoothing makes it difficult to pinpoint the trial-specific contribution of each of the two hypothesized controllers. Research on habits and behavioural automaticity in humans could therefore benefit from a task paradigm where the estimation of the controller balance can be inferred without dependence on the recent trial history, and where action-outcome associations are stable over time.

Although habits, as defined in the animal literature, have been notoriously hard to induce and test in humans (de Wit et al., 2018), considerable progress in the development of task paradigms for the study of habits in humans has recently been made. In one task by Hardwick and colleagues, participants learned to associate four different stimuli with four different responses (pressing one of four keys on the computer keyboard) (Hardwick et al., 2019). One group of participants practiced this stimulus-response association extensively over 4,000 trials across four consecutive days, while a second group only practiced for an average of 40 trials until the S-R associations were learned (Hardwick et al., 2019). Participants then had to learn a revised S-R mapping where the responses of two of the stimuli were switched. Hardwick et al. (2019) showed that when available response times were short, participants in the four-day-training group increased their “habitual” responses, selecting the responses of the now-incorrect but extensively practiced S-R mapping more often than the group without extensive training. Importantly, there was no effect between groups when response times were allowed to be long (>600 ms). Hardwick and colleagues suggested that habits are latently active even in situations where choice behaviour is goal-directed, and that these latent habits can be unmasked when available response times are too short for the slower goal-directed responses to be expressed. Using another experimental approach, Luque et al. (2020) used the reinforcer devaluation as known from the animal literature on habits (Dickinson et al., 1983) and devised a devaluation study for humans. They showed that when using reaction times instead of response choices as a measure of habitization, habitual behaviour can be observed in humans, but only under time pressure, similar to the findings by Hardwick and colleagues. This is consistent with some basic characteristics that are usually associated with habits, namely their capacity for faster and more accurate performance.

To investigate the interaction between habitual and goal-directed behaviour with purely trial-based goaldirected value processing under time pressure, we developed a novel paradigm, which we call the Action-Sequence Task (AST). In the AST, participants implicitly learn an action sequence similar to the so-called serial reaction time task (SRTT) (Robertson, 2007; Lewicki et al., 1988; Nissen and Bullemer, 1987), while occasionally and probabilistically being prompted to act in accordance with an explicitly instructed goal-directed task, in the presence of a demanding time limit. In choice trials of this goal-directed task, participants are asked to quickly choose one of two different response options. In terms of the general task requirement (collecting as many points as possible to maximize monetary payout), these two response options are either equally optimal (i.e. one is as good as the other) or unequal (i.e. one response option should be preferred when acting in a goal-directed manner). While the classical SRTT tests motor sequence learning, the present task in addition requires the learning of action-outcome contingencies, i.e. the probabilities of reward given a key press, and their comparison in dual-target trials. We therefore consider the sequences of key presses as (cued) action sequences, in line with the ideomotor theory of actions, which suggests that actions are bilaterally associated with their anticipated effects (Shin et al., 2010). As reward contingencies stay constant throughout the experiment, the afforded values to the response options in the goal-directed task are trial-based and independent of the recent trial history. Importantly, in such situations participants still have the possibility to act according to the implicitly learned and chunked action sequence, i.e. act habitually (Dezfouli and Balleine, 2012; Balleine and Dezfouli, 2019), which allows us to investigate the interaction of the two different response processes (implicit chunked action sequence vs explicit goal-directed task structure). Our aim was to investigate the interaction of chunked action sequences with goal-directed task requirements, when the habitual response is in conflict with the goal-directed response, when they are in agreement with each other, or when the explicit goal-directed task is uninformative in terms of the response selection process.

## 2 METHODS

### 2.1 Task

The task was performed online and is a modification of the serial reaction time task (Nissen and Bullemer, 1987). In our task participants were shown four boxes on the screen, see Fig. 1. Each of the four boxes were associated with one of four keys on the computer keyboard (s,x,k, and m). In each trial, participants had 600ms to press the key associated with the box in which the target appeared. The positions in which the targets appeared were pseudo-random in the random condition (R), while they appeared in a fixed sequence of twelve alternating target positions in the sequential condition (S, Fig. 1B). The sequence was structured such that it contained all four possible target position three times and did not contain single-target repetitions (the same target appearing twice in a row). Each target was followed by the same target only once. In random blocks, there was no repeating sequence and targets appeared in a pseudo-random order, subject to the constraints that there were no repetitions of the same single-target position, and that two-target cycles repeated twice at most (so e.g. 1-2-1-2-3-… was allowed, but 1-2-1-2-1-… was not allowed). The experiment was conducted over two days, with 14 alternating sequential and random blocks, each containing 480 trials. Sessions lasted around 60 minutes on day one, and 90 minutes on day 2, including instructions.

**Figure 1.**
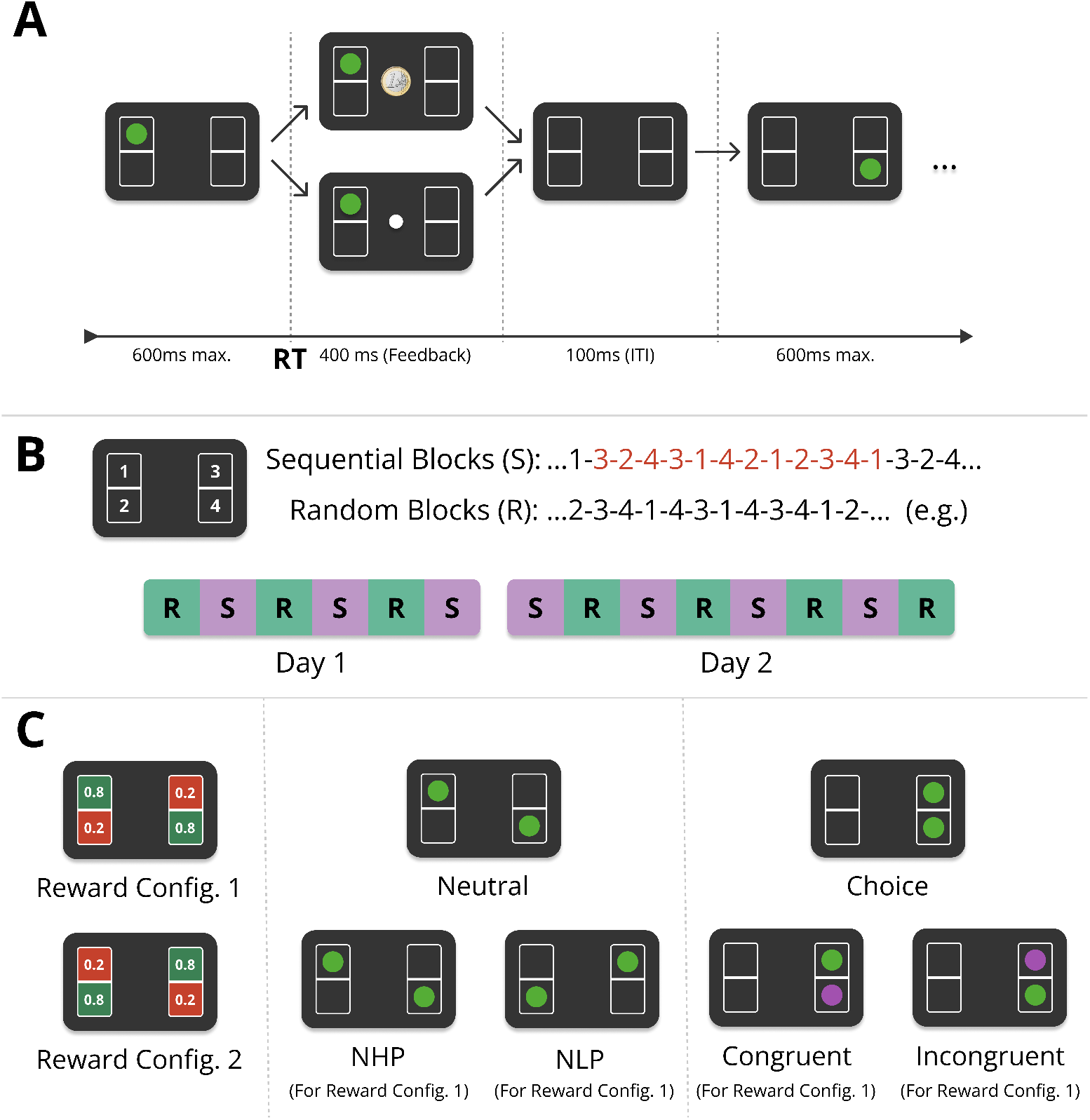
Experimental design of the Action-Sequence Task (AST).(**A**) In each trial, participants saw one or two green target stimuli placed in four different boxes (here a single target is shown). Participants had to press one of four corresponding keys on the keyboard in response to the target position(s), within a deadline of 600ms after stimulus onset. When pressing the correct key, a reward was paid out probabilistically. If the trial was rewarded, a euro coin was shown in the centre of the screen. In the case of no reward, a white dot appeared instead. The feedback (FB) lasted for 400 ms (or 250 ms, depending on the experimental group and phase of the experiment, see Fig. S1). After the feedback phase, there was an intertrial interval (ITI) of 100 ms. (**B**) There were two experimental conditions: In the *sequential* condition (S), targets kept repeating the same twelve-item sequence, while in the *random* condition (R), targets appeared pseudorandomly (red numbers) (see main text for details). The experiment was performed over the course of two consecutive days with alternating sequential and random blocks, six blocks on day one and eight blocks on day two, each block consisting of 480 trials. (**C**) (Left) The reward-probabilities were associated with specific positions on the screen and were either 0.8 or 0.2. Reward-probabilities did not change over the entire experiment and were counterbalanced with one half of participants being presented with reward configuration 1 (RC1), and the other half with RC2. (Center & Right) Dual-target trials (two green targets) appeared pseudo-randomly, with a frequency of between 68 (14.2%) and 78 (16.3%) per block. In the sequential condition, one of the two targets always corresponded to the current sequence element. Both targets can have the same reward-probability (*neutral* (dual-target) trials), Center, or different reward probabilities (*choice* (dual-target) trials, Right). *Neutral trials* are further differentiated into trials where both targets have a low reward probability (neutral low-probability (NLP) trials), or a high probability (neutral high reward-probability (NHP)). *Choice trials*, when appearing in the sequential condition, are further differentiated into two different trial types. If the sequence element (here in purple for illustration, green in experiment) was in a high reward-probability location, it is a *congruent* trial, otherwise it is called an *incongruent* trial.

If participants responded erroneously, i.e. did not respond within 600ms, pressed more than one or an incorrect key, a penalty screen appeared for 1, 200 ms with a text saying “Zu langsam!” (“Too slow!”), “Falsche Taste!” (“Wrong key!”), or (“Bitte drücken Sie nur eine Taste gleichzeitig.”/ “Please press only one key at a time”). If participants responded correctly, they were probabilistically rewarded with a point, indicated by a euro coin appearing on the screen, or a white dot if no point was won. Each box had a probability of either 0.2 or 0.8 to yield a reward in case of a correct response (i.e. no timeout, no double key-press, or wrong key-press) (Fig. 1C left). In each trial, the outcome was sampled randomly based on the corresponding reward probability of the box position. Participants were explicitly instructed which boxes had a higher chance of reward (but not the exact probabilities), that they should choose the response option with the highest reward probability in the case of dual-target trials (see below), and that their final monetary payout increased with the number of points they gained.

To investigate conflict between habitual and goal-directed control, we additionally interspersed dual-target trials (Fig. 1C, center and right panel). Therefore, participants were instructed that sometimes throughout the experiment, two green targets can appear on the screen instead of one (*dual-target trials*, see 2.1.1). For dual-target trials, participants were explicitly instructed to choose the better response option. If the proportion of high reward-probability responses in non-neutral dual-target trials dropped below 65% for a whole block, participants received a feedback at the end of the block reminding them to choose the better response options when they can. Such feedback was conditionally given on the first day, and after the first block on the second day.

#### 2.1.1 Dual-Target Trials

In the sequential condition, a dual-target trial always showed the current sequence element, and a pseudorandomly chosen second target. A dual-target trial never contained the target of the directly preceding single-target trial. Dual-target trials were always followed by two to nine single-target trials, so a dual-target trial was never followed directly by another one. Participants were not told about the two conditions, nor that targets would appear in a fixed order. Each of the fourteen blocks consisted of 480 trials, ≈ 15% of which were dual-target trials (as dual-target trials were positioned pseudo-randomly, the percentage of dual-target trials varies slightly between blocks from 14.2% to 16.3%, while being the same for each participant).

Dual-target trials were classified according to the degree of conflict between habitual and goal-directed control, depending on experimental condition and the two target positions. If a dual-target trial contained one target in a box with a high reward probability, and another target with low reward probability, we will call this a *choice trial* in the following (Fig. 1C), since participants can choose between a response with a high reward probability and a response with a low reward probability. If both targets have the same reward-probability (both low or both high), the trial is designated a *neutral trial*. In the sequential condition, *choice trials* are further categorized into two different types, depending on the reward probability of the target associated with the current sequence element, see Fig. 1D. If the current sequence element of the fixed sequence was located in a box with a high reward probability, in the following we call this a *congruent trial*, as the sequential choice is congruent with the reward-optimal choice. Otherwise, the choice trial is an *incongruent trial*.

#### 2.1.2 Pace-Switch and Counterbalancing

In order to investigate the impact of temporal variations on action sequence chunking and goal-directedness, we introduced a pace-switch condition for half of the participants on the second half of the second day. While the response feedback after every trial was shown for 400 ms throughout the first day and the first half of day two, in the pace-switch group it was shown for 250ms in the second half of day two. The feedback duration remained 400ms for the other group. The pace-switch (PS) group and the group with no pace-switch (NoPS) were further divided into two equally sized groups. Both in the PS and the NoPS groups, half of the participants performed the experiment with reward-configuration 1 (see Fig. 1C), while the other half with reward configuration 2, as well as mirrored stimulus succession for both sequential and random conditions. Contrary to our initial hypothesis of more efficient action sequence chunking in case of reduced inter-trial-intervals in the pace-switch condition, we found no significant effects of the pace-switch. For better legibility, the results can be found in the supplementary material (Fig. S2).

Furthermore, half of the participants saw exactly the same stimulus succession in random and sequential blocks, along with reward-configuration 1 (RC1, see Fig. 1C), while the other half of participants was presented with a stimulus succession that was swapped from the left to the right side (so if one half of the participants saw 3-2-4-3, the other half saw, 1-4-2-1) and the reward-configuration 2 (RC2).

#### 2.1.3 Criterion Test

After the instructions but before the main experiment, participants were told that their understanding of the task will be tested by a short test. In the test, participants were presented 13 successive dual-target trials, ten of which were choice trials. Participants were instructed that there is no time-limit during the test trials, and that they should always choose the option with the higher reward probability when possible. The criterion test failed if a participant chose a low-reward-probability option in more than one choice trial, in which case participants were informed about their failure and again instructed to choose the response option with the high reward-probability when possible. The test could be performed for a maximum number of three times. If a participant failed all three criterion tests, they were excluded from further participation. Of 131 participant who started the experiment, 12 failed this criterion test (fail rate 9.2%).

### 2.2 Participants

Participants were recruited via the central participant pool of the Technische Universität (TU) Dresden. Exclusion criteria were current psychological or psychiatric disorders, age less than 18 or more than 40 years, and current frequent playing of a keyboard instrument. The experiment was hosted online on servers of the TU Dresden center for information services and high performance computing (ZIH) with expfactory (Sochat et al., 2016), and participants performed the experiment online. 131 participants started the experiment. 12 participants failed the criterion test, and 13 were excluded due to miscellaneous problems: 8 had technical problems that resulted in premature discontinuation of the experiment or lost data, one forgot running the experiment on the second day, one discontinued the experiment on day two due to lack of motivation, two participants did not finish the experiment due to reasons that are not traceable (participants did not reply to e-mail), and one participant was excluded because reaction times showed to be consistently below zero, which we ascribe to a technical error. In total, we collected behavioral data of 106 participants. To keep groups balanced with regard to sequence-counterbalancing, we randomly excluded six participants to obtain four groups of equal sizes (*N* = 25 per group) to ensure that the succession of stimuli in the whole experiment is balanced across both hands (for instance, if the sequence element in a specific incongruent trial is in the left hand for one half of participants, it is in the right hand for the other half). Note that the reported results do not qualitatively change when including the six excluded participants. The resulting group of 100 participants consisted of 76 females and 24 males, aged 25.24 *±*4.9 (*μ ± σ*) years, with 91 right-handed, 8 left-handed, and one ambidextrous participant. Data was collected over the course of two months, where data for group 3 were collected one month prior to the other three groups in order to test the internal online data-collection infrastructure. At the end of the experiment on day two, participants were asked (i) whether they had noticed phases in which the experiment appeared easier, (ii) whether they noticed that there was a repeating sequence of twelve keypresses, (iii) whether they could reproduce the twelve-item sequence ad-hoc, (iv) whether they could reproduce the twelve-item sequence after being given the first four keypresses in the sequence, (v) whether they noticed the pace-switch on day two (the last question was only asked to the Pace-Switch group). Ethical approval was granted by the Ethics Committee of the Technische Universität Dresden (EK 514122018).

### 2.3 Data Analysis Strategy

If participants internalized the repeating action sequence in the sequential condition, we expect faster mean reaction times and lower error rates than in the random condition in single-target trials. Furthermore, we expected participants to show a training effect from day one to day two, independent from the task condition, which would manifest in a main effect of day. To test these two key hypotheses of a main effect of task condition (random vs sequential) and a learning effect on reaction times, we performed a two-way repeated measures ANOVA on the participant-specific mean reaction times across all participants with the within-subject factors task condition (sequential (S) and random (R)) and day (day one and day two). We further expected to see similar differences in error rates, with reduced error rates in the sequential condition than in the random condition, as well as a training effect from day one to day two. To test this, we performed the same two-way repeated measures ANOVA as described before, but on the error rates (timeouts, wrong keypresses, or more than one keypress at once).

In an exploratory analysis, we tested whether goal-directed responding as measured by the ratio of high reward-probability choices in choice trials (henceforth called *H*.*P. choice frequency*) was affected by task condition and day of experiment, again by performing a repeated measures ANOVA with the factor task condition (S vs R) and day (day one vs day two). Results indicated this was the case on day two, but not on day one.

To look into this effect in more detail, we investigated the impact of the implicitly learned sequence on the two different types of choice trials, incongruent and congruent trials, on day two vs on day one. Our third key hypothesis was that of a differential effect on choice-trial type (increased H.P. choice frequency in congruent trials and decreased H.P. choice frequency in incongruent trials). We performed a two-way repeated measures ANOVA on the H.P. choice frequencies in sequential choice trials with the first factor being the dual-target type (congruent vs incongruent) and the second factor the experimental day (day one vs day two). Results aligned with our hypothesis of a differential effect of trial-type on H.P. choice frequency. We further found that the impact of the repeating sequence on choice behaviour increased from day one to day two.

If choice behaviour between congruent and incongruent trials is more affected by the automatic action sequence on day two, we expected to see larger differences in reaction times between congruent and incongruent dual-target trials on day two (congruency effect). We tested this in an exploratory analysis, comparing reaction times between congruent and incongruent trials in each sequential block of the experiment.

We then hypothesized that, if the action sequence is learned, it should manifest as faster reaction times in sequential blocks. At the same time, a stronger effect of the action sequence should manifest in choice behaviour as an increased H.P. choice frequency in congruent trials and a decreased H.P. choice frequency in incongruent trials. We therefore expected that reaction times in the sequential condition should correlate with the difference in H.P. choice frequencies of congruent and incongruent trials, which we tested with Pearson correlation.

Lastly, in an exploratory analysis, we tested for a main effect of dual-target type on reaction times. We performed a one-way repeated measures ANOVA on the mean reaction times with the single factor being the dual-target type. We performed this ANOVA with four levels (incongruent, congruent, NLP, NHP) for the sequential condition, and three levels in the random condition (choice, NLP, NHP). To test a similar effect of dual-target type on error rates, we performed a two-way repeated measures ANOVA for each condition (sequence and random), with the first factor being the dual-target type, and the second factor the error type (Timeouts vs Other Errors (being wrong keypresses or double keypresses)). We differentiate between timeout errors and other errors because the analysis on reaction times showed that participants have increased reaction times for NLP trials, and reaction times above 600 ms would be considered a timeout, confounding the effect of long reaction times with other error types. To investigate the effect of the automatic action sequence on choice behaviour in the different dual-target types, we tested whether sequential responding in the sequential condition was different from the chance level of 50% for the four different dual-target types using two-tailed single-sample t-tests for each dual-target type (congruent, incongruent, NLP, and NHP). Lastly, we compared the effect of the learned action sequence between pairs of the four dual-target types in the sequential condition with paired t-tests on the mean sequential choice frequencies (i.e. ratio of sequential responses to all responses in dual-target trials in the sequential condition).

### 2.4 Code and Data Availability

The experimental paradigm with which data collection was done, raw data, and analysis code are available on the open science framework (https://osf.io/dsb4a/). DOI: 10.17605/OSF.IO/DSB4A.

## 3 RESULTS

### 3.1 Participants internalize action sequence

We tested for an effect of task condition and time on reaction times, using a repeated measures ANOVA with factors task condition (S vs R) and day (day one vs day two). We found significant main effects of both task condition (*F* (1, 99) = 262.51, *p* < 0.0001, partial *η*^2^ = 0.04) and day (*F* (1, 99) = 72.7, *p* < 0.0001, partial *η*^2^ = 0.08) (see Fig. 2A). For error rates, a repeated measures ANOVA yielded a significant main effect of task condition (*F* (1, 99) = 119.4, *p* < 0.0001, partial *η*^2^ = 0.04) and a significant main effect of day (*F* (1, 99) = 29.46, *p* < 0.0001, partial *η*^2^ = 0.03), as well as a small but significant interaction effect between task condition and day (*F* (1, 99) = 8.65, *p* = 0.004, partial *η*^2^ = 0.003). Participants on average responded faster while at the same time making less errors in the sequential condition than in the random condition. This shows that participants internalized the action sequence and exploited their implicit knowledge about the sequence for faster and less erroneous responses. The main effects of day shows that participants on average get faster from day one to day two, and perform less errors overall. The interaction effect shows that the effect of day is not the same for the two condition types. The speed-up of response times from day one to day two is stronger for the sequential condition than for the random condition.

**Figure 2.**
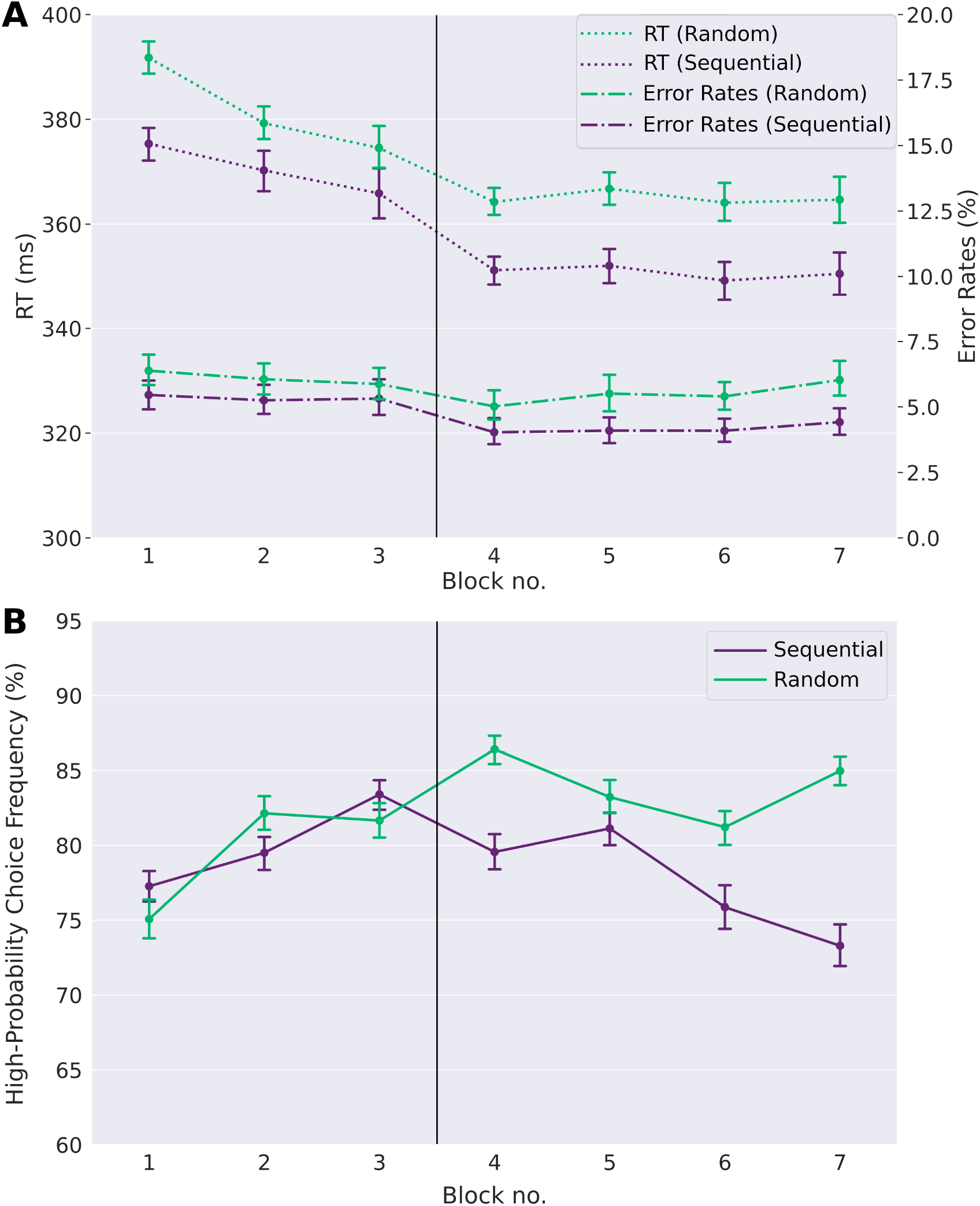
Learning of automatic action sequence and goal-directed task.(**A**) Reaction times (RT) and error rates (ER) (timeouts and wrong responses) in the random and the sequential condition across single-target trials. Participants show faster reaction times and lower error rates in the sequential condition than in the random condition, evidenced by a main effect of task condition (see text). This is already the case in the first blocks of day one, suggesting that participants quickly internalized the repeating sequence to a certain degree, exploiting the anticipation of upcoming target positions for fast reaction times. Furthermore, participants’ reaction times decrease over time, as do the error rates, showing a learning effect with time. (**B**) high reward-probability (H.P.) choice frequencies in choice trials. H.P. choice frequencies remain far above the chance level of 50% in both experimental conditions, showing that participants generally acted in a goal-directed manner. However, while there is no difference between the experimental conditions on day one, a difference emerges on day two. As can be seen, high reward-probability choice frequencies on day two are reduced in the sequential condition compared to the random condition. Error-bars show standard errors of the means.

Note that error rates and response times are smaller already in the first sequential block compared to the first random block. While this might be due in part to the sequence learning effect, this is likely also the result of a cue-response training effect in the first random block, where participants get acquainted with the general task structure, since all participants started with a random block on day one.

### 3.2 Participants choose goal-directed actions

Next, we tested whether participants were also able to act in agreement with the goal-directed task. As can be seen in Fig. 2B, responses in choice trials were generally goal-directed in both conditions, as the ratio of high reward-probability choices in dual target trials is well above chance level of 50% throughout the whole experiment. Across all blocks and participants, choice frequencies for the high reward-probability option in choice trials were on average 80.3% (ranging between 56.1% and 96.5%). When testing for the effects of task condition and day using a repeated measures ANOVA, we found a main effect of task condition (*F* (1, 99) = 60.0, *p* < 0.0001, partial *η*^2^ = 0.03), no main effect of day (*F* (1, 99) = 1.5, *p* = 0.23, partial *η*^2^ = 0.002), and a significant interaction between task condition and day (*F* (1, 99) = 93.7, *p* < 0.0001, partial *η*^2^ = 0.04). A closer inspection shows that, while there is no effect of the task condition on high reward-probability choice frequencies on day one (*p* = 0.45, two-sided paired t-test), the effect is significant on day two (*p* < 0.0001, Cohen’s d = 0.64, paired t-test) and clearly visible in Fig. 2B. Participants made fewer high reward-probability choices in the sequence compared to the random condition (77.5% vs. 84.0% on day two, *p* < 0.0001, Cohen’s d = 0.64 two-tailed paired t-test).

### 3.3 Increase of Sequence Impact on Day Two

We next looked at the influence of the implicitly learned sequence on goal-directed actions in congruent and incongruent dual-target trials, so-called choice trials, see Fig. 1C. If the reduced high reward-probability choice frequencies in the sequential condition on day two (see Fig. 2B) are due to the effect of the learned sequence, we expected to see a differential effect in the choice behaviour of congruent and incongruent trials.

A repeated measures ANOVA on the high reward-probability choice frequencies yielded a significant main effect of choice trial type (*F* (1, 99) = 77.8, *p* < 0.0001, partial *η*^2^ = 0.12), a significant main effect of day (*F* (1, 99) = 194.6, *p* < 0.0001, partial *η*^2^ = 0.36), and a significant interaction between these two factors (*F* (1, 99) = 77.8, *p* < 0.0001, partial *η*^2^ = 0.12). This main effect of choice trial type can be readily seen in Fig. 3, where participants generally chose the response option with the high reward-probability more often in congruent than in incongruent trials. This reflects the relative increase of this choice difference from day one to day two. In random blocks, in the absence of a repeating action sequence, the frequency of high reward-probability choices increases from day one to day two and can be interpreted as a baseline without sequence influence. On both days, participants choose high reward-probability response more often in congruent trials than in random choice trials (85.5% in congruent trials, 79.6% in random choice trials on day one, *p* < 0.0001, Cohen’s d = 0.62 day one, 90.7% in congruent trials and 84.0% in choice trials on day two, *p* < 0.0001, Cohen’s d = 0.86, two-tailed paired t-tests) and less often in incongruent trials than in random choice trials (75.0% in incongruent trials on day one, *p* < 0.0001, Cohen’s d = −0.39, and 68.1% on day two, *p* < 0.0001, Cohen’s d = 1.19, two-tailed paired t-tests). Participants seem to learn the action sequence quickly, with a difference in the high reward-probability choices between congruent and incongruent trials of 25.9% in the second half of the first sequential block, compared to 10.6% to the first half, and no difference within the first 40 trials of the first block.

**Figure 3.**
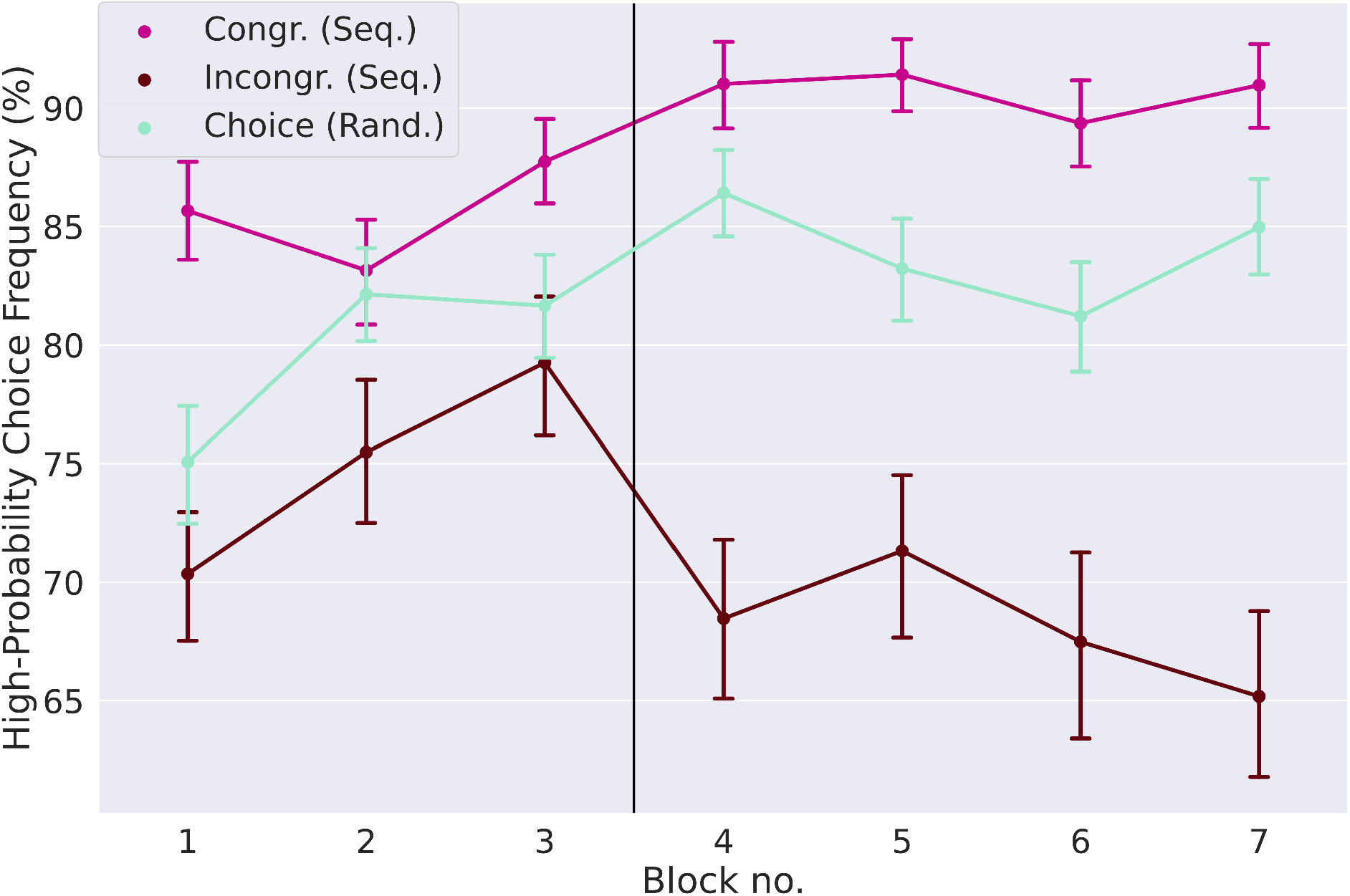
Influence of implicitly learned sequence on goal-directed action. The high reward-probability (H.P.) choice frequencies (a measure of goal-directed task performance, see methods) are shown for the three different types of choice trials: Congruent and incongruent trials in the sequential condition, and choice trials in the random condition. In incongruent trials the H.P. choice frequency is lower relative to the random condition, while it is higher in congruent trials. This shows that the preference for a sequential response not only creates a conflict in incongruent trials, but also aids the decision process in congruent trials. The distance between the H.P. choice frequencies for congruent and incongruent trials, which is a measure of the strength of the learned action sequence, increases from strongly from day one to day two. Error-bars show standard errors of the means.

We expected that reaction times are faster in congruent trials than in incongruent trials (congruency effect, Fig. 4). Slower reaction times in incongruent trials indicate that the response conflict between the two task modalities (goal-directed and automatic) increases. Although the effect is rather small, it is larger than zero in sequential blocks four and seven on day two (*p* < 0.0001, Cohen’s d = 0.35 block four, Cohen’s d = 0.37 block two, two-sided t-tests, p-values Bonferroni-corrected for seven comparisons).

**Figure 4.**
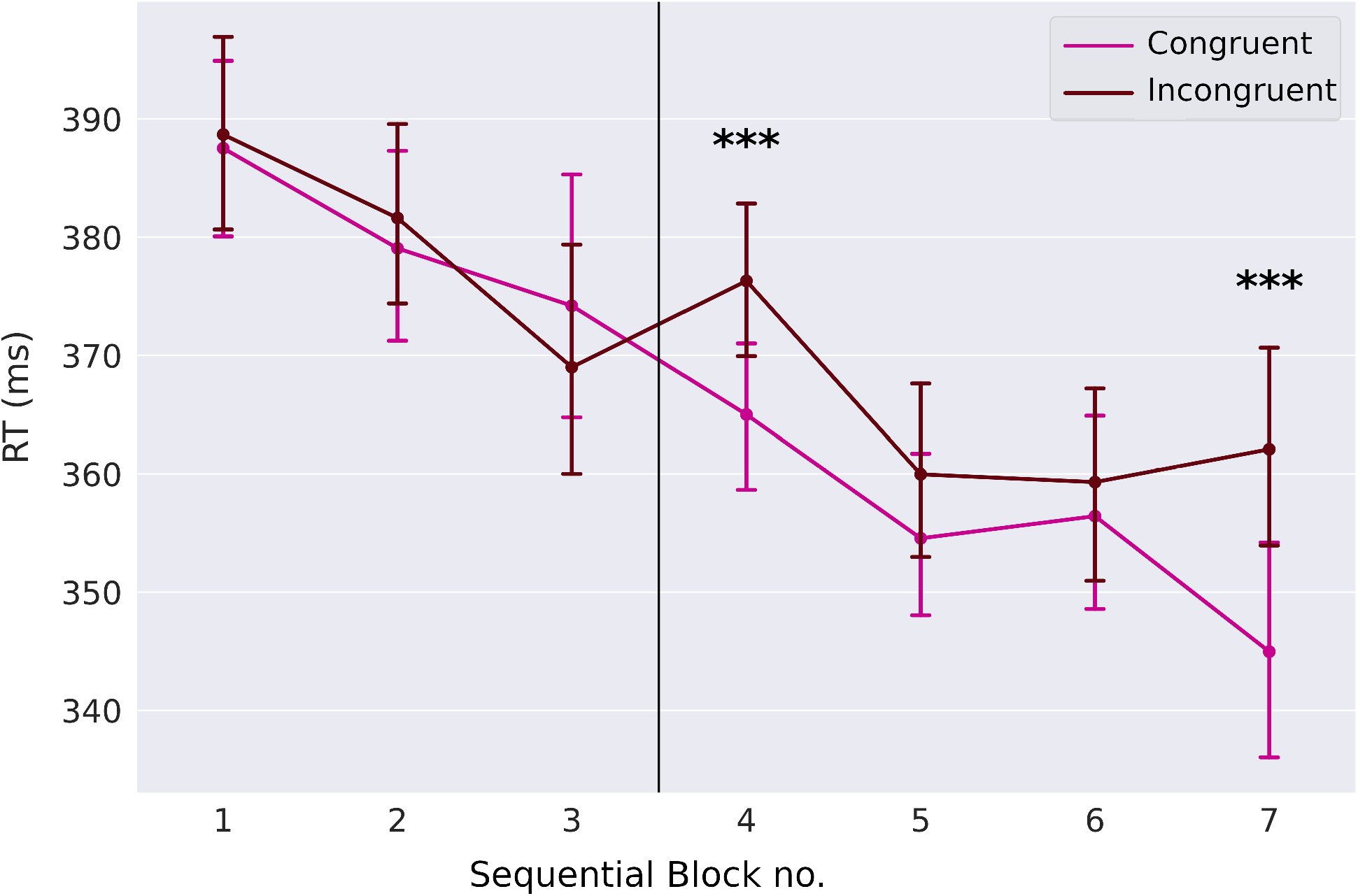
Congruency effect. While reaction times for congruent and incongruent trials are not different on day one, they are different on blocks four and seven on day two. Error-bars show standard errors of the means.

We also observed a significant Pearson correlation between individual reaction time differences for single-target trials between sequential and random condition, and the choice difference of high reward-probability choice frequencies between congruent and incongruent trials on both days (*r* = 0.37, *p* < 0.001 (Day 1), and *r* = 0.54, *p* < 0.0001 (Day 2), Fig. 5). We further observed a similar relationship between the H.P. choice difference and the participants’ individual error rates between sequential condition and random condition in single-target trials (*r* = 0.24, *p* = 0.02 (Day 1), and *r* = 0.43, *p* < 0.0001 (Day 2)). In both cases the correlation is increased significantly from day one to day two. These results give further support to an influence of the chunked action sequence on choice behaviour, as well as to its effect on reaction times and error rates.

**Figure 5.**
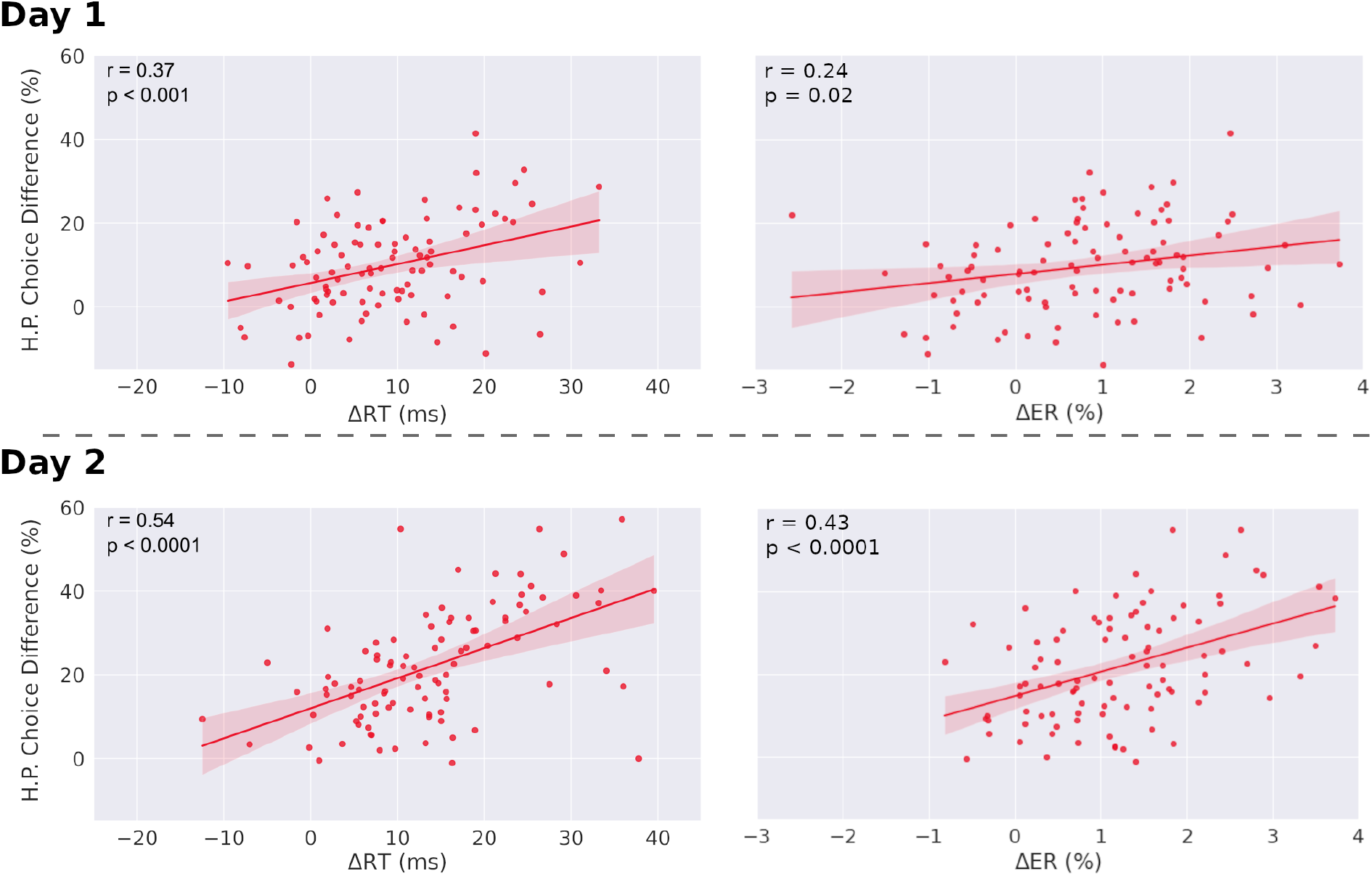
Correlations between choice differences and both reaction time differences and error rate differences, in single-target trials. Left: Correlation of individual H.P. choice frequency for congruent and incongruent trials (H.P. choice difference) and mean reaction time differences between sequential and random condition for single-target trials. Both slopes are significantly different from zero. The correlation coefficient of day two is significantly increased relative to day one (*p* < 0.0001). Right: Correlation of individual H.P. choice frequency for congruent and incongruent trials (H.P. choice difference) and mean error rate differences between sequential and random condition for single-target trials. Both regression slopes are significantly different from zero. The correlation coefficient of day two is significantly increased relative to day one for both error rates and reaction times (*p* < 0.0001). The regression slope further increases from day one to day two for error rates (*p* = 0.018), but not for reaction times (*p* = 0.1). Δ*RT* = *RT*_Random_ − *RT*_Seq_, Δ*ER* = *ER*_Random_ − *ER*_Seq_. Analysis was performed after outlier removal (outliers were defined as elements more than three scaled median absolute deviations from the median). Results before outlier removal are similar for reaction times. For error rates, before outlier removal, the correlation is not significant for day one (*r* = 0.16, *p* = 0.12), but significant for day two (*r* = 0.40, *p* < 0.0001).

### 3.4 Neutral trials

In neutral trials, in order to investigate the effect of dual-target type on reaction times, we performed a one-way repeated measures ANOVA with the factor trial type for both experimental conditions (Fig. 6A). Note that for neutral trials, all effects over time, akin to Fig. Fig. 2 and Fig. 3, can be found in the supplementary material (Fig. S3-S6), along with the results of paired t-tests (Tables S1-S3). In the sequential condition we found a significant main effect of trial type (*F* (3, 297) = 292.1, *p* < 0.0001, partial *η*^2^ = 0.27), with significant pairwise differences between congruent and incongruent trials (*p* < 0.0001, Cohen’s d =− 0.15 two-tailed paired t-test), as well as between NLP trials and all other dual-target types (NLP vs congruent *p* < 0.0001, Cohen’s d = 1.34; NLP vs incongruent *p* < 0.0001, Cohen’s d = 1.23; NLP vs NHP *p* < 0.0001, Cohen’s d = 1.26; two-sided paired t-tests, p-values are Bonferroni-corrected for five comparisons). While mean reaction times for congruent, incongruent and NHP trials all range between 365 ms and 371 ms, mean reaction times for NLP trials are 415 *±* 38 ms. In the random condition, we also found a significant main effect of trial type (*F* (2, 198) = 407.7, *p* < 0.0001, partial *η*^2^ = 0.98). We further found significant pairwise differences between all three pairs of dual-target trial-types (choice vs NHP *p* < 0.0001, Cohen’s d = − 0.14; NLP vs choice *p* < 0.0001, Cohen’s d = 1.37; NLP vs NHP *p* < 0.0001, Cohen’s d = 1.18; two-sided paired t-tests with Bonferroni correction for three comparisons). Here, mean reaction times for choice and NHP trials are 382 ms and 387 ms, while for NLP trials mean reaction times are 429 ms.

**Figure 6.**
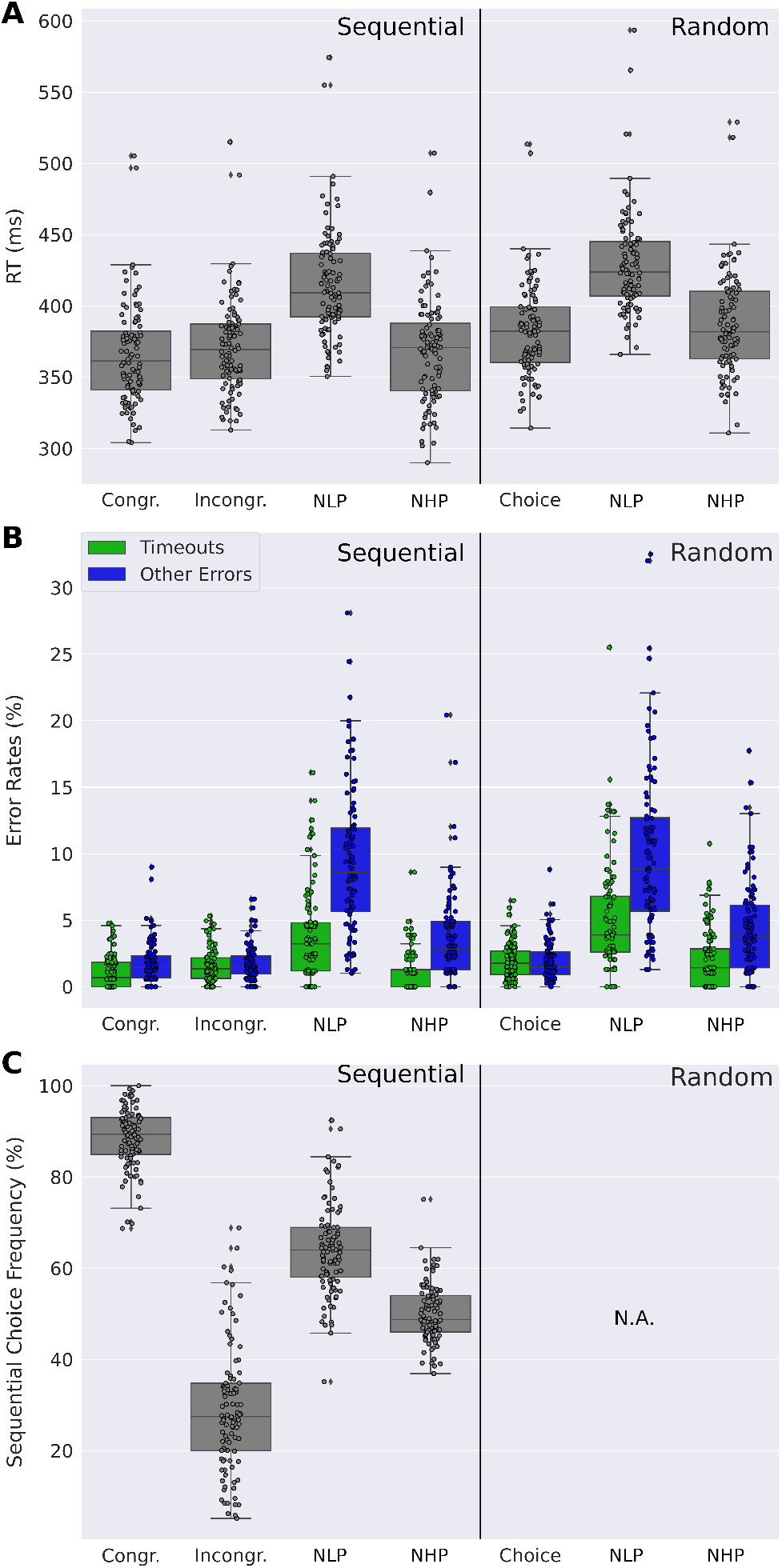
Comparison of dual-target trials.(**A**) Distribution of participants’ mean reaction times for different types of dual-target trials: Choice trials (Congruent (Congr) and Incongruent (Incongr) in the sequential condition), Neutral Low Reward-Probability (NLP) and Neutral High Reward-Probability (NHP) trials. Participants show markedly increased reaction times for NLP trials. (**B**) Distribution of participants’ mean number of timeouts and of other error types (i.e. wrong or double key-presses). Timeouts and other errors are elevated for NLP trials relative to other dual-target trials, in both the sequential and random conditions. Errors other than timeouts are increased also for NHP trials compared to other non-neutral dual-target trial types in both conditions. ((**C**)) Distribution of participants’ mean sequential choice frequency for different dual-target trials. While sequential choice frequencies are at chance level for NHP trials, they are at 64.2% and above chance level for NLP trials. **A**-**C** Data were averaged across both days. Dots show the corresponding mean values of the individual participants.

To test whether dual-target types similarly affect errors, we performed a two-way repeated measures ANOVA on dual-trial type and error type for each experimental condition (Fig. 6B). In the sequential condition we found a significant main effect of trial type (*F* (3, 297) = 170.5, *p* < 0.0001, partial *η*^2^ = 0.71), a significant main effect of error type (*F* (1, 99) = 83.2, *p* < 0.0001, partial *η*^2^ = 0.42), and a significant interaction between trial type and error type (*F* (3, 297) = 65.3, *p* < 0.0001, partial *η*^2^ = 0.39). We further found significant pairwise differences between NLP trials and all other trial types, for both error types (timeouts and other), all with medium to large effect sizes (all *p* < 0.0001, two-tailed paired t-tests, Bonferroni corrected for 14 comparisons, Cohen’s d between 0.9 and 1.9. For the full results, see Tables S1-S3). In the random condition we further found a large significant main effect of trialtype (*F* (2, 198) = 170.6, *p* < 0.0001, partial *η*^2^ = 0.61), a large significant main effect of error type (*F* (1, 99) = 45.49, *p* < 0.0001, partial *η*^2^ = 0.29), and a large significant interaction between trial-type and error type (*F* (2, 198) = 36.5, *p* < 0.0001, partial *η*^2^ = 0.22). Significant pairwise differences were also found between NLP trials and both other trial types, for both error types, with medium to large effect sizes (all *p* < 0.0001, two-tailed paired t-tests, Bonferroni corrected for nine comparisons, Cohen’s d between 0.9 and 1.7).

Finally, we analyzed the sequential choice frequencies in the four different dual-target trials (congruent, incongruent, NLP, and NHP) (Fig. 6C). As can already be seen in Fig. 3, sequential responding is strongly different between congruent and incongruent trials. Interestingly, sequential responding is different from the chance level for all dual-target types (*p* < 0.0001, two-tailed t-tests, Bonferroni corrected for four comparisons), except for NHP trials (*p* = 0.9, uncorrected). Furthermore, in NLP trials, sequential choice frequency (64.2%) is significantly higher than in NHP trials (50.1%; *p* < 0.0001, Cohen’s d = 1.7, two-tailed paired t-test). Increased sequential choice frequencies in NLP trials, relative to NHP trials, were observed in every sequential block throughout the whole experiment, ranging from a difference of 7.9 (in sequential bock 6) percentage points to 19.6 percentage points (in sequential block 2). The effect cannot be explained by a preference for a certain hand (for example the dominant hand), as the positions of the sequence elements in NLP and in NHP trials were equally distributed between the right and the left hand across participants. It is however interesting to note that participants have an increased tendency to respond with their dominant hand in NLP trials (61.0% *±* 9.5%) compared to NHP trials (53.9% *±* 22.4%; *p* < 0.0001, Cohen’s d = 0.34, two-tailed paired t-test). Significance tests were two-tailed t-tests signed-rank tests to test whether sequential choice frequencies were different from the chance level of 50%.

In summary, we found that in non-neutral sequential choice trials (congruent and incongruent dual-target trials), participants behaved as expected: Participants in general chose the high reward-probability target in dual-target trials with high probability, but with more training the learned sequential response preference interfered with this task (Fig. 3). However surprisingly, in neutral trials, we observed a much reduced effect of the sequential choice preference (Fig. 6C). For neutral high reward-probability target trials, this difference was most striking with a sequential choice frequency at chance level, i.e. 50%.

## 4 DISCUSSION

Here we presented the action sequence task, a paradigm similar to the serial reaction time task (SRTT), but with the crucial extension of another task dimension. This second task dimension consists of explicit task goals, and requires the evaluation and comparison between two response options, therefore requiring goal-directed cognitive processes such as value retrieval and value comparison. Crucially, the goal-directed, explicit task goals were constructed to be sometimes in agreement and sometimes in conflict with the implicitly learned action sequence, therefore allowing for the analysis of the interaction between a chunked action sequence and an explicit goal-directed task.

We found that participants learned and internalized the repeating action sequence as evidenced, in the sequential condition, by both reduced reaction times and high reward-probability choice frequencies (Fig. 2), while still generally acting in accordance with the goal-directed task (Fig. 3). We further found, according to our initial hypothesis, that in congruent trials (i.e. trials where the action sequence and the goal-directed task require the same response) the implicit action sequence increased optimal choices as required by the goal-directed task and therefore helped participants improve performance (Fig. 3). In incongruent trials, goal-directed performance was reduced due to sequential responding. These results show that the learned action sequence interferes with advantageous goal-directed behavior in dual-target trials. Unexpectedly, in neutral trials, we found that the impact of the learned action sequence was low (Fig. 6C). This is surprising, since a greater influence of the action sequence could here be exploited for shorter reaction times (since one admissible response is always the sequential response), with no negative impact on performance, as opposed to incongruent trials, where the influence of the action sequence negatively impacts performance. As the goal-directed task provides the same expected reward to both response options in both neutral trial types, we expected a marked influence of the implicitly learned sequence on choice behaviour to resolve the conflict between the two response options. However, we found the opposite: the impact of the implicit sequential response option was low, especially in the case of neutral trials with high reward-probability reward options (NHP).

How can the obtained results be explained? First, it is evident that participants learned both action preferences afforded by the two different task components, the implicit action sequence and the goal-directed value comparison task component. They performed well above chance-level in choice trials (Fig. 2B), even in incongruent trials, indicating a general tendency for goal-directed responding. Furthermore, participants quickly acquired a chunked action sequence, which is shown by reduced reaction times and error rates in the sequential relative to the random condition (Fig. 2A).

We found that the two response preferences, sequential and goal-directed, interact in congruent and incongruent trials in the sequential condition (Fig. 2B). However, this interaction looks markedly different in neutral trials, where participants’ choices do not show a sequential response preference in neutral high reward-probability (NHP) trials, or show only little interaction in neutral low-probability (NLP) trials, see (Fig. 6C). It is furthermore striking that reaction times and error rates are markedly increased in trials where there is a conflict within the goal-directed system (i.e. in neutral trials where the conflict arises due to two equally good, or bad, response options), but not so much when the conflict arises between the two response systems (i.e. in incongruent trials where the goal-directed system contradicts the automatic system). What is the explanation for these small effects of the sequential choice preference in neutral trials? The reason cannot be that participants simply followed their instructions of choosing a response with high reward probability, or that participants quickly identified a neutral trial to adjust their response preference at the start of a 600 ms trial, because participants did not do this in incongruent trials, either.

Given the high proportion of single-target trials (∼ 85%), it is reasonable to assume that in the sequential condition, participants find the sequential preference highly useful to perform well in the single-target and congruent double-target trials and therefore always have their sequential preference ’switched on’, also in neutral trials. This makes the results on neutral trials surprising, because if both response options in an NLP trial are equally bad, why not resolve this conflict quickly by employing the sequential choice? Rather, there is only a relatively small influence of the sequential preference and, critically, an increased error rate and time-outs, relative to all other dual-target trials, Fig. 6B. An explanation might be that, in NLP trials, only the goal-directed system first tries to resolve the conflict between two equally bad targets. Only later, when the system realizes that time is running out, other options are pursued, even pressing the wrong key (error rate of close to 10%, see Fig. 6B) or, finally, let the sequential preference determine the response. As is evident from the elevated average reaction times and time-out rates, relative to all other dual-target trials, this switching away from a resolution of the conflict within the goal-directed system takes its time. We speculate that a similar process takes place in NHP trials. Here, there is again a conflict only in the goal-directed system, i.e. there is a choice to make between two equally good response options. This conflict can be resolved within the goal-directed system, and switching to the automatic system is not necessary to resolve the conflict, as evidenced by an indifference to the sequential response option. We believe that this explanation for the overall behaviour in dual-target trials can provide insight in the interaction between a hypothetical automatic and a goal-directed system, as described in dual-process theories. Our findings suggest that, under time pressure, it depends on the type of conflict, whether there is an influence of the automatic system on the goal-directed system, or not. Intuitively, if there is a conflict within the goal-directed system, one might assume that the automatic system might be most useful to resolve that conflict, as in our task. However, according to our results this is not what happens under time pressure. Rather, the goal-directed system first tries to resolve the conflict on its own, and only later, when there is the risk of response time running out, breaks off its attempt at a resolution and involves the automatic system to find a response.

In this study, we have further shown that faster reaction times in single-target trials in the sequential condition are associated with increased habitual behaviour in dual-target trials of the sequential condition. We have thus shown that positive measures of habits in the form of faster reaction times and smaller error rates can be measured with the present task. This is important, since many studies on habitual behaviour employ paradigms where habitual behaviour is pitted against goal-directed behaviour, and thus effectively interpret the absence of goal-directedness as habitual behaviour, instead of measuring markers of habitual behaviour per se (Sjoerds et al., 2013; Hardwick et al., 2019; Balleine and Dezfouli, 2019). However, habits are often acquired mostly implicitly, and in the absence of any strongly opposing goal-directed intentions. In our everyday lives, a habit is usually only noticed after it has been acquired, and when it then comes into conflict with an explicit goal, as is the case in so-called slips of actions (Mylopoulos, 2022). For habit research, it is therefore desirable to find a way of measuring habitual behavior as a sui generis mode of action, which can be achieved by measuring positive characteristics of habits independently of goal-directed behaviour, in the form of, for instance, reduced reaction times and error rates.

In addition to measuring the acquisition of habitual behaviour per se, the present study also offers insight into the conflict between goal-directed and habitual behaviour and their arbitration, by means of so-called dual-target trials, where both modes of behaviour, habitual and goal-directed, can be employed to select a response. Future research directions could focus on the dynamic allocation of control within the arbitration process by introducing uninterrupted chains of congruent or incongruent dual-target trials. In such a situation, the dynamic allocation of control in the arbitration process might change towards a reduced impact of the chunked action sequence. The analysis of individual differences in dynamic control allocation might be interesting especially in combination with other established experiments that measure the conflict between goal-directed behaviour and other modes of behaviour, like for instance the Simon task (Simon and Rudell, 1967), where goal-directed control is measured against spatial priming, as opposed to the current study, where response priming is a result of the chunked action sequence. Habits further play an important role in the study of psychopathologies, as it has been theorized that some mental disorders, such as, among others, substance use disorder, obsessive-compulsive disorder, and Tourette syndrome, are associated with a maladaptive reliance of, or a pathological increase in the strength of, habitual behaviour (Graybiel and Rauch, 2000; Vandaele and Janak, 2018; Everitt and Robbins, 2005; Everitt et al., 2016; Ersche et al., 2016; Sjoerds et al., 2013). The investigation of changes in the present measures that occur in mental disorders could therefore help gaining insight into clinically relevant changes in the reliance of habits in behaviour selection.

While one class of habit theory viewed habits as stimulus-response associations that are acquired through reinforcement, this appears to be at odds with key aspects of habitual behaviour that seem intuitive from everyday experience. For example, some habits as tying shoes or washing hands are often performed in the same way, but do not usually result in any obvious reinforcing reward. Furthermore, such complex types of habits which consist of a sequence of individual actions do not figure well with the idea that a habit is a rather simple action in response to a triggering stimulus, unless the complex action sequence like washing hands is itself considered the habit. Consequently, new models of habitual behaviour have emerged which incorporate these aspects of habits. In (Miller et al., 2019), the authors propose a habit model where single-action habits are formed through mere repetition. In this model, habits can arise in the absence of reinforcement and habits are balanced against a goal-directed controller which evaluates actions based on model-based planning. While the model by Miller and colleagues considers habits as single actions, (Dezfouli and Balleine, 2012, 2013), propose a hierarchical model of action selection, where habits are modelled by action sequences which are chunked into so-called macro actions, and are activated by a supraordinate goal-directed controller. This controller evaluates the value of a goal-directed action and its alternative, the habitual macro action. Importantly, macro actions can be performed faster than goal-directed actions and can therefore maximize obtained reward per unit of time. Although we did not design our experiment in such a hierarchical fashion, there may be an interesting parallel to this hierarchical view of Dezfouli and Balleine. In our study, the expected ongoing execution of an action sequence is interrupted by the dual-target trials. In a hierarchical view, this interruption of the habitual action sequence and switching to another task might be handled by a higher-level controller. While this is originally not covered by the model proposed by Dezfouli and Balleine, where a macro action, once chosen, is executed until termination, the idea of hierarchical, goal-directed action control with selection of habitual action sequences is an important consideration in contrast to the flat view of balancing between habitual and goal-directed control. While the hierarchical model proposed by Dezfouli and Balleine has already been shown to reproduce experimental data of the two-stage task (Dezfouli and Balleine, 2013), it will be interesting to assess whether it can replicate key aspects of the present study. In a third approach, (Schwöbel et al., 2021) proposed a hierarchical Bayesian model that combines the idea of habit acquisition through repetition and habits as action sequences. In this model, habits are considered as precise priors over action sequences in a Bayesian integrator model, where the value-based goal-directed mode of behaviour is represented by a Bayesian likelihood function. Prior and likelihood are combined to compute a posterior over actions from which an action is chosen. The model was shown to reproduce key findings from habit literature in rodents (Schwöbel et al., 2021). The model is explicitly based on an inference of context, where different contexts are identified by their reward structure and state-transitions. Each context is associated with a different prior over action sequences, which resonates with the hierarchical view of (Dezfouli and Balleine, 2012). Taken together, one view of our results may be to assume that a controller at a higher level switches between the two tasks (single- and dual-target trials). In this view, the conflict would not be directly between habitual and goal-directed actions but be at the task level. We will further investigate this possibility in future work.

Note that the interpretation of the present results are limited by two aspects of the present experiment. The first limitation is that it is unclear whether the reinforcer in the form of a point reward (indicated by a coin on the screen) is a necessary or sufficient motivation for participants to guide their actions. In pilot studies preceding the experiment presented here with different task parameters, we noticed that participants often ignored the reward-probabilities of different response options in dual-target trials, which was in part due to the demanding deadline, as was noted by some participants. This changed after we introduced the criterion test along with detailed instructions, reminding participants to choose the better response option in dual-target trials when they could, and feedback about their high reward-probability choice frequency at the end of each block (see Sec. 2.1). While the achieved effect was the same to a reward-driven motivation (namely, participants developed a goal-directed preference for two of the four response options), future research will have to investigate whether reinforcer feedback is in fact necessary after every trial, and how different motivators affect the experimental outcome.

To date, most of habit theory stems from animal research, and only few experiments in humans have succeeded in observing classical measures of habitual behaviour derived from studies in rodents (Hardwick et al., 2019; Luque et al., 2020; Tricomi et al., 2009). In order to learn more about habitual and goal-directed behaviour in humans, we need an assay of experiments that can reliably measure positive measures of habitual behaviour, and investigate the interaction between habits and goal-directed behaviour in a lab setting. As we have shown here, such studies might be most useful under tight deadline regimes, which is exactly where fast habitual behaviour is most needed, similar to our dynamic everyday environment which often requires fast responses due to the interaction with conspecifics. Our study hints at the possibility that tight deadlines are not exclusively the realm of automatic fast responses, but that there may be an intricate interaction pattern between habitual and goal-directed system that produces fast and flexible behaviour, even under time pressure.

## AUTHOR CONTRIBUTIONS

SF, ME and SJK designed the study. SF and ME implemented the study, collected and analysed the data. SF wrote most parts of the article. SF, ME, SJK, TE, and MNS interpreted the data. SJK, TE, and MNS contributed to the article and all authors approved the submitted version.

