## Supplementary Material for "Interaction between habits as action sequences and goal-directed behavior under time pressure"

### 1 PACE-SWITCH

In order to investigate the impact of short deadlines on automatic sequential behaviour, we introduced a pace-switch condition at the end of the experiment, reducing the response-to-stimulus intervals for one group of participants at the end of day two, while the other group did not experience such a reduction S1A. Participants were not told about the pace-switch. We hypothesized that this pace-switch adds to the time pressure, increasing further a detectable impact of the automatic system on the decision-making process, leading to increased sequential responding, as task-switching experiments have frequently shown that goal-directed responding is often impeded by shorter response-to-stimulus intervals (Monsell, 2003). We therefore compared sequential responding in dual-target trials of the pace-switch group to sequential responding in the non-switch group (Fig. S2). We found no group-differences in sequential responses in incongruent and congruent trials ($p > 0.5$ (uncorr.), two-tailed paired t-tests for congruent and incongruent trials). Surprisingly, we found that sequential responding in the first sequential block after the pace-switch is lower in the pace-switch group, for neutral trials (Fig S2) ($\Delta = 6.1$ p.p., $p = 0.02$ (uncorr.), two-tailed paired t-test; however this difference is not significant after a Bonferroni correction for the three different dual-target trial types). When investigating the pace-switch effect on other behavioural measures (reaction times, timeout rates, error rates, HP choice frequency), we found no differences between the two groups.

We therefore conclude that the pace-switch did not lead to changes in participants' response behaviour.

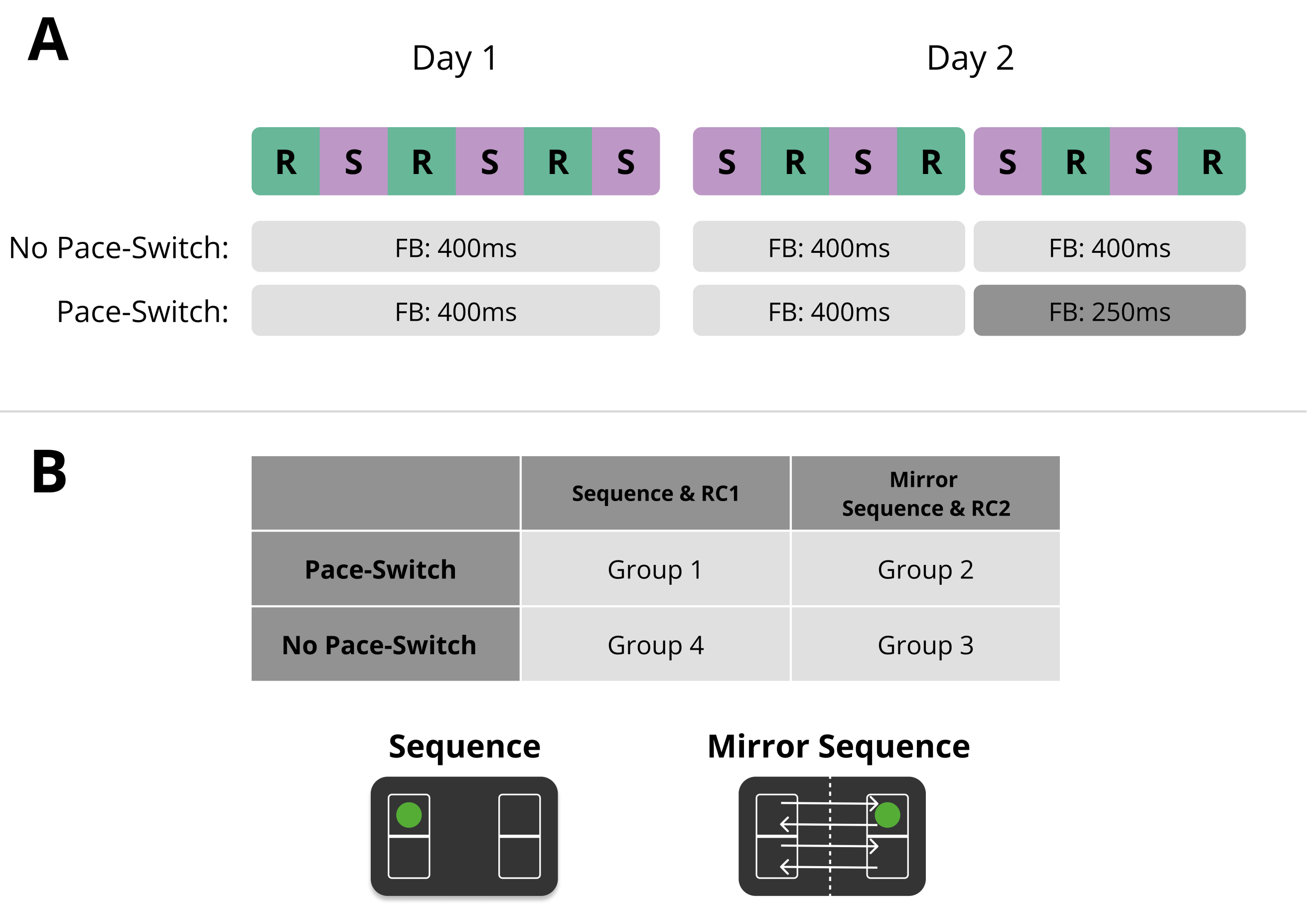


**Figure S1.** (**A**) During the last four blocks of day two, the feedback duration (Paper Fig. 1) for half of the participants ($n = 50$) was shortened from 400 ms to 250 ms. (**B**) The experiment employed a 22 factorial design with one factor for the Pace-Switch. Regarding the other factor, one half of the participants saw the identical stimulus succession across the whole experiment, along with reward configuration 1 (RC1). The other half saw a mirrored version of the stimulus sequence, along with RC2. Notice that the term "sequence" here refers to the stimulus succession of the whole experiment (including random blocks). Through mirroring stimulus succession and reward configuration at the same time, dual-target trials kept their type. Of 50 participants who saw the mirrored sequence, 45 were right-handed, 4 left-handed, 1 ambidextrous. Of 50 participants who saw the other sequence, 47 were right-handed and 3 left-handed.

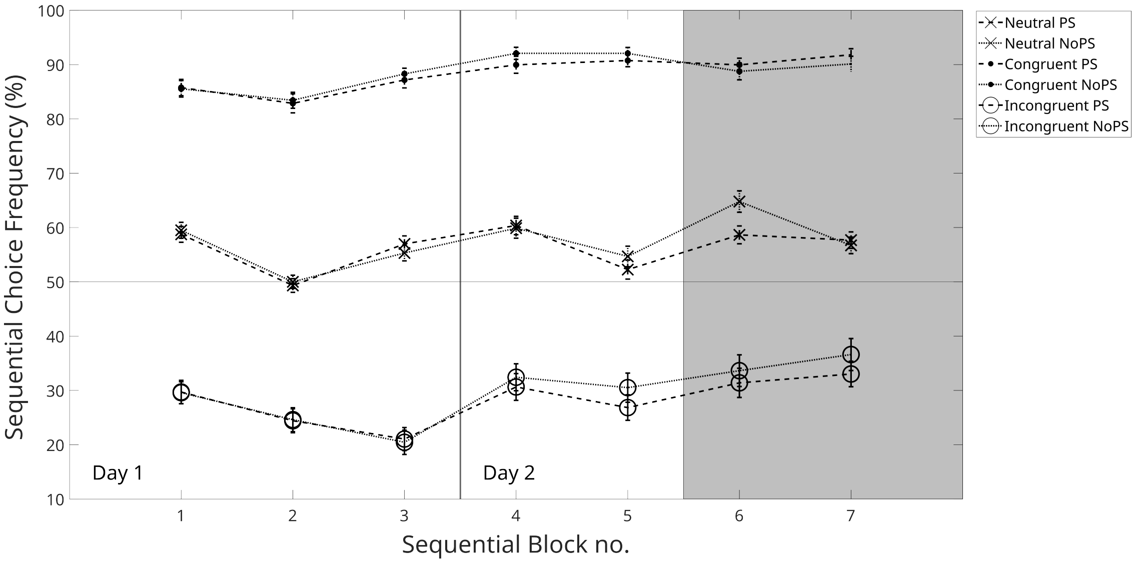


**Figure S2.** Sequential choice frequency for dual-target trials. Here we show the ratio of sequential responses to all responses in dual-target trials in the sequential condition, for different dual-target trial types. The gray-shaded area indicates the phase of the experiment where half of the participants experienced a pace-switch (PS) and the other half no pace-switch (noPS). Throughout the whole experiment, sequential choice preferences were the same for the pace-switch group as for the no pace-switch group. In the sequential block directly following the pace-switch there is a significant difference in sequential choice preference between the pace-switch condition and the no pace-switch group, but the significance is not robust to a multiple-comparisons correction for three comparisons ($p = 0.02$, uncorrected). Furthermore, contrary to our initial hypothesis, the group who did not experience a pace-switch showed an increased sequential choice preference in the first sequential block after the pace-switch. Error bars show standard errors of the mean.

### 2 NEUTRAL TRIALS

**Table S1.** Uncorrected p-values of two-tailed paired t-tests and effect sizes for pairwise comparisons of reaction times.

| Comparison | p-value (uncorr.) | Cohen's d |
|---|---|---|
| Sequential Condition | | |
| Congr vs Incongr | $3.5 \cdot 10^{-6}$ | -0.15 |
| Congr vs NLP | $3.3 \cdot 10^{-41}$ | -1.34 |
| Congr vs NHP | 0.17 | -0.05 |
| Incongr vs NLP | $1.2 \cdot 10^{-34}$ | -1.23 |
| Incongr vs NHP | 0.07 | 0.09 |
| NLP vs NHP | $2.0 \cdot 10^{-33}$ | 1.26 |
| Random Condition | | |
| Choice vs NLP | $9.2 \cdot 10^{-42}$ | -1.37 |
| Choice vs NHP | $3.6 \cdot 10^{-5}$ | -0.14 |
| NLP vs NHP | $2.6 \cdot 10^{-36}$ | 1.18 |

**Table S2.** Uncorrected p-values of two-tailed paired t-tests and effect sizes for pairwise comparisons of errors.

| Comparison | p-value (uncorr.) | Cohen's d |
|---|---|---|
| Sequential Condition — Timeout Rates | | |
| Congr vs Incongr | 0.0028 | -0.33 |
| Congr vs NLP | $3.5 \cdot 10^{-14}$ | -1.07 |
| Congr vs NHP | 0.13 | 0.20 |
| Incongr vs NLP | $7.5 \cdot 10^{-11}$ | -0.90 |
| Incongr vs NHP | $4.2 \cdot 10^{-6}$ | 0.49 |
| NLP vs NHP | $7.5 \cdot 10^{-15}$ | 1.14 |
| Sequential Condition — Other Errors | | |
| Congr vs Incongr | 0.71 | 0.04 |
| Congr vs NLP | $7.0 \cdot 10^{-26}$ | -1.87 |
| Congr vs NHP | $2.4 \cdot 10^{-9}$ | -0.77 |
| Incongr vs NLP | $4.2.5 \cdot 10^{-27}$ | -1.91 |
| Incongr vs NHP | $7.3 \cdot 10^{-10}$ | -0.81 |
| NLP vs NHP | $8.1 \cdot 10^{-18}$ | 1.22 |
| Random Condition — Timeout Rates | | |
| Choice vs NLP | $1.4 \cdot 10^{-13}$ | -1.03 |
| Choice vs NHP | 0.44 | -0.09 |
| NLP vs NHP | $1.7 \cdot 10^{-12}$ | 0.91 |
| Random Condition — Other Errors | | |
| Choice vs NLP | $2.0 \cdot 10^{-24}$ | -1.74 |
| Choice vs NHP | $3.5 \cdot 10^{-12}$ | -0.89 |
| NLP vs NHP | $1.2 \cdot 10^{-15}$ | 1.10 |

**Table S3.** Uncorrected p-values of two-tailed paired t-tests and effect sizes for pairwise comparisons of sequential choice frequencies.

| Comparison | p-value (uncorr.) | Cohen's d |
|---|---|---|
| Congr vs Incongr | $7.9 \cdot 10^{-55}$ | 5.48 |
| Congr vs NLP | $2.3 \cdot 10^{-37}$ | 2.91 |
| Congr vs NHP | $2.3 \cdot 10^{-70}$ | 5.95 |
| Incongr vs NLP | $2.1 \cdot 10^{-53}$ | -2.92 |
| Incongr vs NHP | $4.6 \cdot 10^{-26}$ | -1.95 |
| NLP vs NHP | $4.6 \cdot 10^{-23}$ | 1.70 |

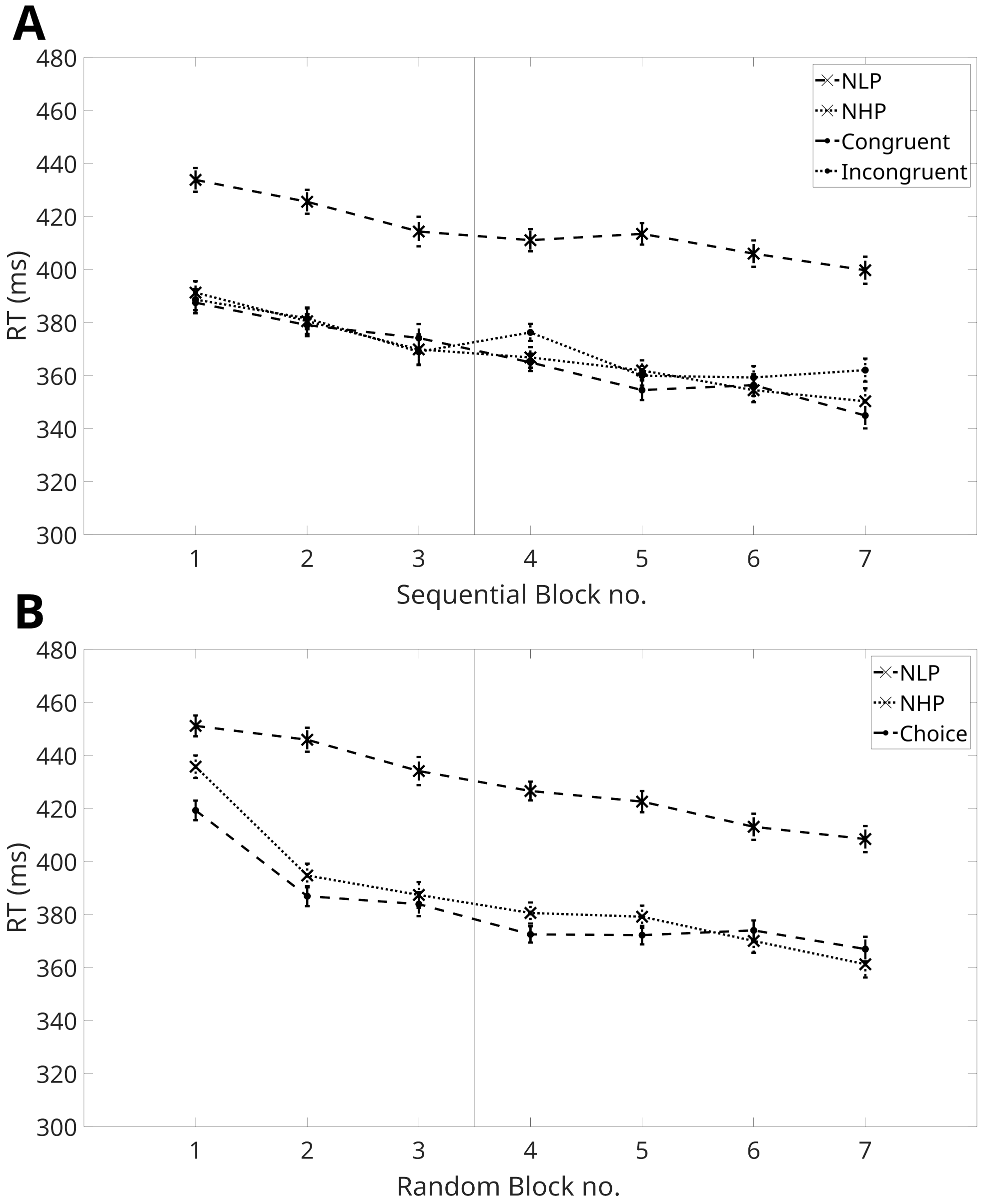


**Figure S3.** **A** Reaction times for dual-target trial types in the sequential condition. **B** Reaction times for dual-target trial types in the random condition. Error bars show standard errors of the mean.

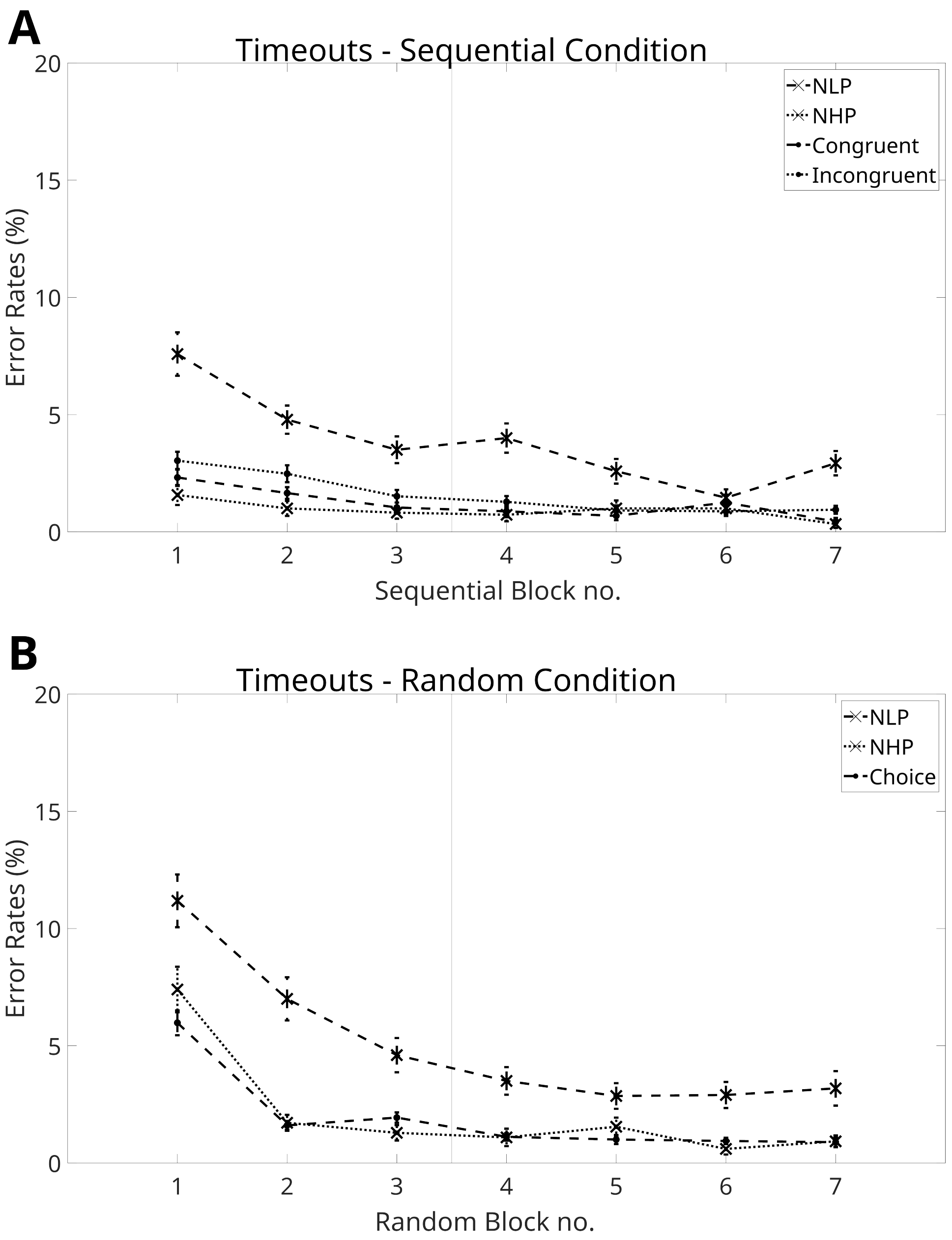


**Figure S4.** **A** Timeout rates for dual-target trial types in the sequential condition. **B** Timeout rates for dual-target trial types in the random condition. Error bars show standard errors of the mean.

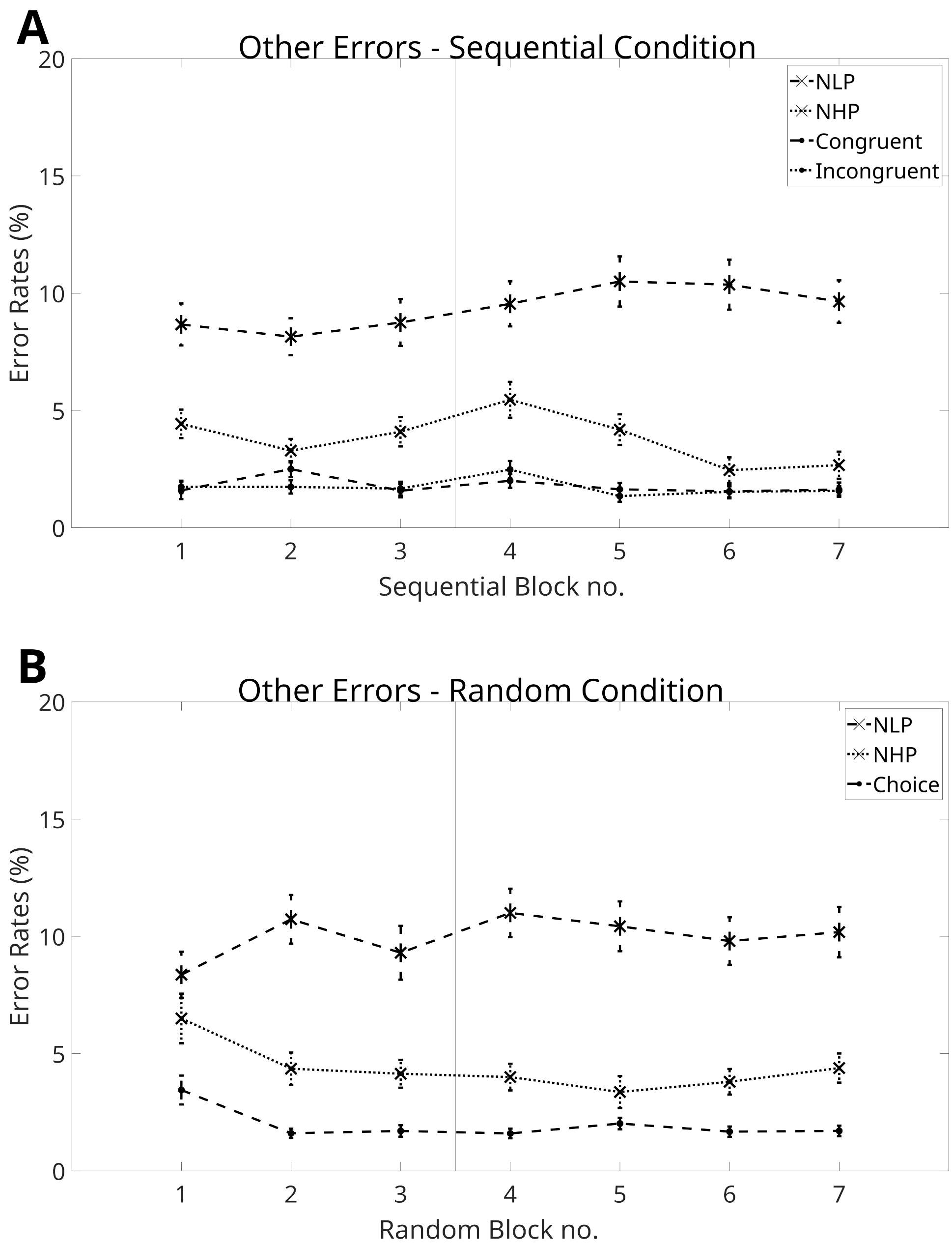


**Figure S5.** (**A**) Rates for errors other than timeouts for dual-target trial types in the sequential condition (**B**) Rates for errors other than timeouts for dual-target trial types in the random condition. Error bars show standard errors of the mean.

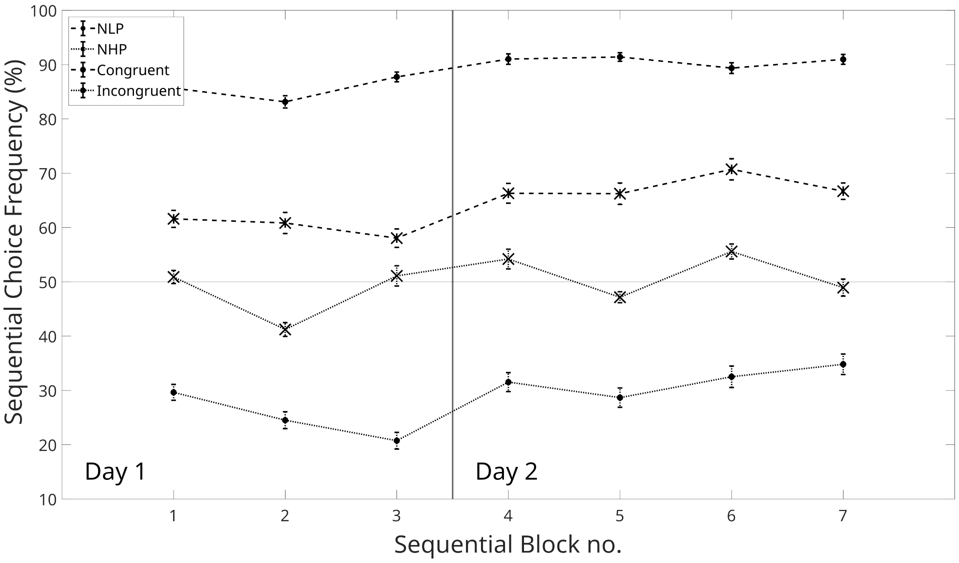


**Figure S6.** Sequential choice frequency for dual-target trial types in the sequential condition. Error bars show standard errors of the mean.

### 3 POST-EXPERIMENT QUESTIONNAIRE

Question 1: “Did you have the impression that over some phases of the experiment, you found it easier to make less or no mistakes?”
Results: yes 87%, no 8%, Don’t know 5%.
Question 2: “Throughout the experiment, a sequence of twelve key presses was often repeated. Did you notice this?”
Results: yes 33%, no 57%, Don’t know 10%.
Question 3: “Please try to repeat the previously mentioned sequence, or parts of it, by pressing the keys s,x,k, and m in the corresponding order.”
Results: On average, participants were able to reproduce 20.6% of the sequence. This was measured as the longest contiguous sequence of keys in the participant’s answer, that occurred as a subsequence in the actual sequence. 28% of participants did not reproduce any subsequence. One participant was able to reproduce the whole sequence.
(Only PS Group) Question 4: “In the second part of today’s experiment, the pace of appearance of the green dots changed. Did you notice this?”
Results: yes 68%, no 24%, Don’t know 8%.
